# Collection and Storage of HLA NGS Genotyping Data for the 17^th^ International HLA and Immunogenetics Workshop

**DOI:** 10.1101/116871

**Authors:** Chia-Jung Chang, Kazutoyo Osoegawa, Robert P. Milius, Martin Maiers, Wenzhong Xiao, Marcelo Fernandez-Viňa, Steven J. Mack

**Author notes:** Corresponding Author: Steven J. Mack, Center for Genetics, Children’s Hospital Oakland Research Institute Oakland, CA 94609, USA.

## Abstract

For over 50 years, the International HLA and Immunogenetics Workshops (IHIW) have advanced the fields of histocompatibility and immunogenetics (H&I) via community sharing of technology, experience and reagents, and the establishment of ongoing collaborative projects. In the fall of 2017, the 17^th^ IHIW will focus on the application of next generation sequencing (NGS) technologies for clinical and research goals in the H&I fields. NGS technologies have the potential to allow dramatic insights and advances in these fields, but the scope and sheer quantity of data associated with NGS raise challenges for their analysis, collection, exchange and storage. The 17 ^th^ IHIW has adopted a centralized approach to these issues, and we have been developing the tools, services and systems to create an effective system for capturing and managing these NGS data. We have worked with NGS platform and software developers to define a set of distinct but equivalent NGS typing reports that record NGS data in a uniform fashion. The 17th IHIW database applies our standards, tools and services to collect, validate and store those structured, multi-platform data in an automated fashion. We are creating community resources to enable exploration of the vast store of curated sequence and allele-name data in the IPD-IMGT/HLA Database, with the goal of creating a long-term community resource that integrates these curated data with new NGS sequence and polymorphism data, for advanced analyses and applications.

**Abbreviations:**

**Abbreviations**
CSV: Comma-Separated Values
GFE: Gene Feature Enumeration
GL: Genotype List
HLA: Human Leukocyte Antigen
HML: Histoimmunogenetics Markup Language
H&I: Histocompatibility and Immunogenetics
IHIW: International HLA and Immunogenetics Workshop
IMGT: ImMunoGeneTics
IPD: ImmunoPolymorphism Database
IUPAC: International Union of Pure and Applied Chemistry
KIR: Killer-cell Immunoglobulin-like Receptor
MIRING: Minimum Information for Reporting Immunogenomic NGS Genotyping
NGS: Next Generation Sequencing
PI: Principal Investigator
RSCA: Reference Strand Conformation Analysis
rSSO: Reverse Sequence-Specific Oligo
SBT: Sequence-Based Typing
sFTP: secure File Transfer Protocol
SS: Sequence-Specific
SSO: Sequence-Specific Oligo
SSP: Sequence-Specific Priming
WMDA: World Marrow Donor Association
WS: Workshop
XML: eXtensible Markup Language

## 1. Introduction

### 1.1 The Histocompatibility Workshops

Since their introduction in 1964, the Histocompatibility Workshops have been forums for the exchange of community knowledge and experience, allowing histocompatibility and immunogenetics (H&I) researchers, clinicians and technologists to evaluate new methods and technologies, establish standards and advance ongoing collaborative projects. Sixteen International HLA and Immunogenetics Workshop (IHIW) meetings have been held on five continents over the last half-century[1–16], and the 17th IHIW will be held in northern California in the fall of 2017, continuing many long-standing workshop projects.

The advent of next-generation sequencing (NGS) based genotyping technologies has allowed new insights and innovations for the fields of histocompatibility, immunogenetics and immunogenomics. The ultimate goals of the 17th IHIW are to advance H&I basic research and clinical efforts through the application and evaluation of NGS HLA and KIR genotyping technologies, and to foster the development of NGS technologies tailored to meet the H&I community#x2019;s needs, building on the technological and scientific momentum of the previous sixteen workshops.

Toward these ends, we have developed systems, standards and tools for the collection, storage and management of NGS HLA genotyping data (the HLA genotype and associated consensus sequences) generated for 17th IHIW projects. The goals of this effort are to build on the data-collection and -storage experiences of previous workshops, and produce NGS data-managing tools that will support IHIW efforts and persist as public resources after the 17th IHIW. Here, we provide a brief overview of the challenges faced organizing coordinated data-generation and -collection efforts, the strategies we have applied, and the tools, standards and services we have developed to address these challenges.

### 1.2 The Challenges of Coordinated Data Collection

The collection, storage and analysis of data have been key issues of all workshops. Many workshops have used centralized databases[17–21], while in several of the more recent workshops, individual components and projects were responsible for collecting, managing and analyzing data[22–32]. Centralized data-management requires close communication between workshop participants and leaders, instrument and software vendors, and database developers to achieve consensus regarding required data content, data formats, reporting guidelines and quality standards. Sufficient time is also required for all parties involved to develop both the systems and tools to manage data, and the preliminary data on which to test the tools.

#### 1.2.1 Reference Data Management

The specifics of the H&I field bring additional challenges that any data-management and analysis approach, centralized or decentralized, must address[33]. The body of HLA sequence data and associated allele names curated by the IPD-IMGT/HLA Database[34] (Reference Database) increases every four months; because workshop data-generation efforts often span multiple years, the details of the pertinent Reference Database version under which each HLA genotype was generated must be collected along with the genotyping data. The collection and management of genotyping meta-data such as these (described in Table 1) can be just as important for the workshop effort as the genotyping data themselves; without them it may not be possible to determine the extent to which datasets generated years apart or using different methods are equivalent. When workshop efforts span time periods that include major changes to the nomenclature[35, 36], these problems are only compounded.

**Table 1.**
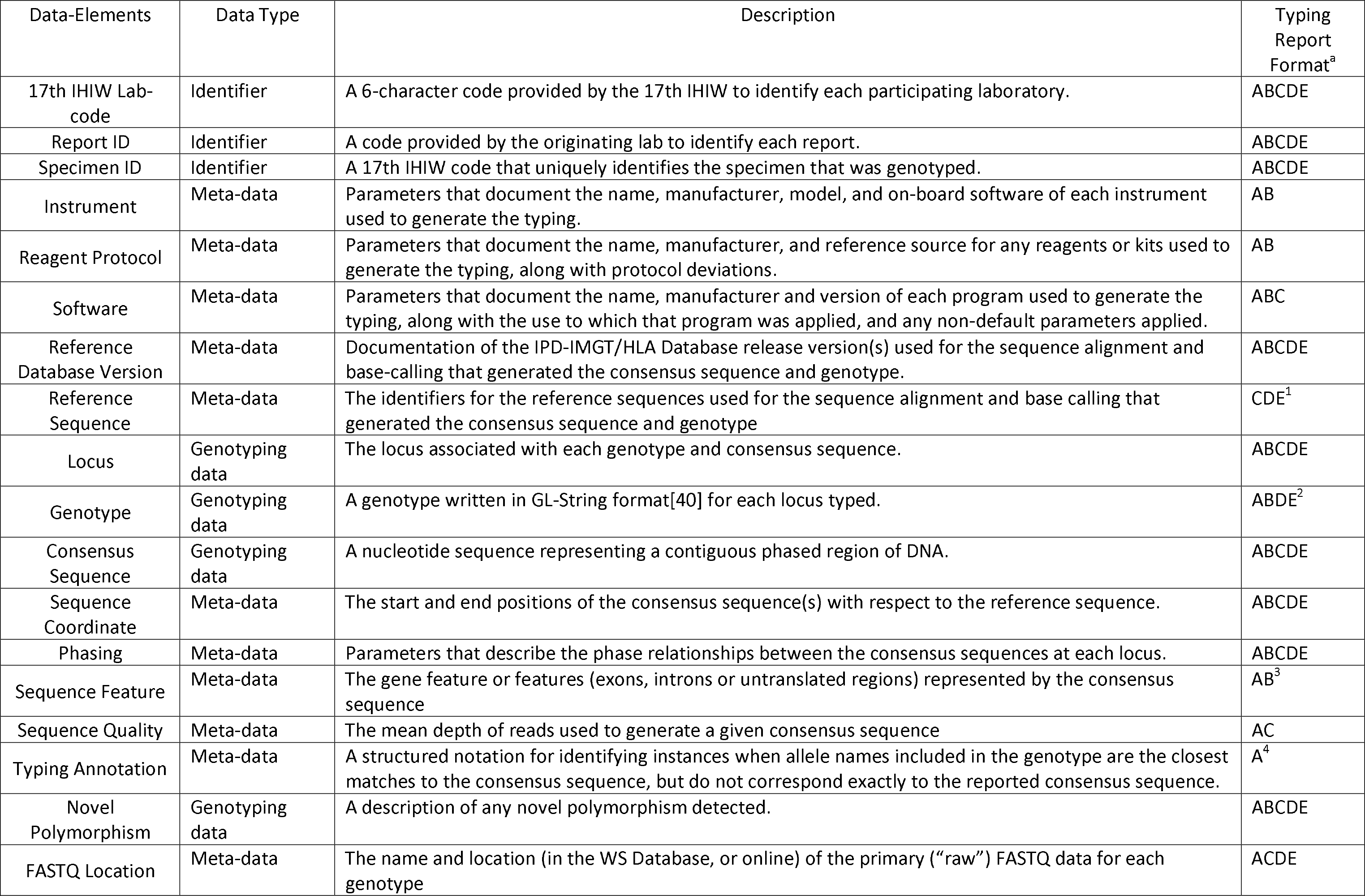
Data-Elements in 17th IHIW Typing Reports.

#### 1.2.2 Primary Data Management

The nature of the primary or “raw” data, from which all experimental data and meta-data are ultimately derived, can vary widely from method to method and from project to project. This was particularly pronounced for the molecular genotyping methods applied in the 11^th^ through the 16 ^th^ workshops, where multiple reference strand conformation analysis (RSCA), sequence-specific (SS) oligo (SSO), reverse SSO (rSSO), SS priming (SSP) and sequence-based typing (SBT) methods were in use, each with its own distinct type of primary data.

#### 1.2.3 Allele Name Data Management

Allele name data must be recorded and managed in a standard manner to facilitate meaningful data-analysis. For many of the previous workshops, the management of HLA allele names has been performed by humans, and involved data recorded in paper documents or spreadsheets in a variety of different ways. Humans are adept at “figuring out” the true meaning of unusual notations and spreadsheet-initiated errors that may occur, but machines are not. For example, “HLA-A*02:99” and “HLA-A*03:01:02” are often recorded as “02:99” or “03:01:02” in spreadsheet columns labeled “HLA-A”, “A”, etc.; however, common spreadsheet applications may change “02:99” to “0.152083333333333” or “3:39”, and “03:01:02” to “3:01:02”, all of which erroneously represent times instead of alleles. The range of potential human-generated transcription errors is large. Previous workshops devoted considerable manual effort to review, identify and correct errors, and standardize allele-name notations prior to analysis. However, the analysis, collection, exchange and storage of NGS genotyping data requires machines (computers) that are able to process allele name data, and the accompanying nucleotide sequence data, without the human ability to identify and correct errors.

#### 1.2.4 Describing Novel Polymorphism

The description of previously unknown (novel) HLA sequence variants has been a long-standing challenge for the H&I community. Until a novel sequence is assigned a name by the World Health Organization Nomenclature Committee for Factors of the HLA System (Nomenclature Committee)[37], it is very difficult to discuss that sequence in the context of the HLA nomenclature. The common practice, associated with pre-NGS genotyping, has been to append a “novel-allele” identifier to a truncated version of a related allele name (e.g. “HLA-A*02V”, “HLA-A*02:NEW”, “HLA-A*02:01new”, etc.). The World Marrow Donor Association guidelines for the use of HLA nomenclature (WMDA guidelines) indicate that “NEW” should be reported for alleles that have not been named by the Nomenclature Committee[38]. However, the absence of a standard for describing novel HLA alleles and associated nucleotide sequences represents a considerable challenge for the collection of NGS HLA genotyping data.

## 2. Meeting the Challenge

The 17th IHIW has adopted a centralized data-storage approach, in which all specimen-related data, reference data, genotyping data and associated meta-data are stored in a single database system. The goal of this effort is to facilitate data and analysis access for workshop participants, with these workshop products and the database itself made available to the H&I community after the workshop’s close. The 17th IHIW focus on NGS provides a large advantage for centralized data collection in that there are currently only a small number NGS platforms, which generate primary data in the same format (FASTQ[39]), and associated genotyping software. A key goal for the 17th IHIW is to collect machine-generated HLA data for consumption by IHIW informatics services, with minimal human intervention. We have worked with NGS software developers to develop a small number of equivalent and interchangeable data reporting formats that allow genotyping data and meta-data to be collected using a “uniform NGS data-collection” approach. This approach builds on the work already accomplished developing the genotype list (GL) string format[40] and the GL Service[41], the Minimum Information for Reporting Immunogenomic NGS Genotyping (MIRING) reporting guidelines and messaging standard[42], and the MIRING-compliant Histoimmunogenetics Markup Language (HML) version 1.0 messaging format[43].

### 2.1 Uniform NGS Data Collection

The 17th IHIW does not require that all workshop projects or participating laboratories use the same NGS platform, typing kit or protocol. NGS instruments manufactured by Illumina (e.g., MiSeq), One Lambda (e.g., S5XL), Pacific Biosciences (e.g., PacBio RSII) and Roche 454 (e.g., GS FLX) have been used in 17th IHIW NGS genotyping efforts. The goal in uniform NGS data collection is that all NGS genotyping data and associated meta-data (which together constitute a “typing report”) should be compatible and comparable, so that all collected data are equally interpretable, regardless of the format in which those data are exchanged. This will allow data generated by different laboratories, in different countries, using different platforms and software, to be stored in one database and made available for multiple projects.

Toward this end, the 17th IHIW accepts NGS genotyping data and meta-data in three MIRING-compliant eXtensible Markup Language (XML)[44]-based typing report document formats – HML (version 1.0.1); GenDx XML, exported by GenDx NGS Engine version 2.4.0; and IHIW XML^A^, a format developed specifically for the 17th IHIW (detailed in Supplements A and B). HML is generated by HistoGenetics, Omixon HLA Twin (version 1.1.4.2), Immucor MIA FORA (version 3.1) and One Lambda TypeStream Visual (version 1.1) software. IHIW XML typing reports can be generated using the 17th IHIW Database (WS Database) system (described in section 2.2), by an individual laboratory (using the Supplementary Materials), and by Illumina, using a “.cgp” file exported by TruSight HLA Assign version 2.1 RUO. We are working with Pacific Biosciences to determine the appropriate typing report document format for data generated on PacBio instruments. In addition, the WS Database accepts HLA genotypes in a comma-separated values (CSV) file generated by Scisco Genetics.

GenDx XML, HML and IHIW XML typing reports include subsets of the NGS genotyping data and meta-data elements described in Table 1. These data-elements are equivalent to MIRING elements 1-8[42]. An HML or GenDx XML typing report might include additional data, but because these document formats include equivalent 17th IHIW data-elements, all submitted HML and GenDx XML HLA typing reports can be converted into IHIW XML typing reports (as described in section 2.2.1), which are then stored in the WS Database. In addition to these typing reports, the primary FASTQ data, too large to include in a report, are stored on a secure File Transfer Protocol (sFTP) server linked to the WS Database.

### 2.2 17th IHIW Database

The WS Database includes an Oracle SQL database (12c Standard Edition) and a web application built with APEX 5.0, running on a multi-core Linux CentOS 6 platform with 960GB of storage, expandable up to 3TB. The 17th IHIW sFTP server is an IBM high-performance computing cluster running Linux RedHat 6, with a 1Gbps Ethernet connection. The server comprises a management node, three compute nodes, two storage nodes and 15 TB of storage. The WS Database schema is illustrated in Supplementary Figure S1. WS Database tools and services are scripted in the Perl, R or Python programming languages. Both the database and the sFTP server are housed in the high-performance computing Stanford Data Center facility on the Stanford Linear Accelerator Center campus, and are managed by the Stanford Research Computing staff.

The WS Database’s structure reflects the workshop’s organization and the defined roles of workshop participants. Each of the six 17th IHIW components – NGS of HLA, NGS of KIR, Hematopoietic Cell Transplantation, Mapping of Serologic Epitopes, Informatics of Genomic Data, and Quality Control & Quality Assurance – is led by a Component Chair (or Chairs). Projects are associated with each component, with a Principal Investigator (PI) for each project. PIs can enroll Lab Members, and can enroll in Components and Projects. Lab Members upload and manage data, and enroll as Project and Component Affiliates. Further details of these participant roles can be found online^B^.

The WS Database system^C^ stores data from typing reports and Scisco CSV tables, FASTQ files, subject and specimen data, and pedigrees (PED format[45]), and manages the accounts and data-access privileges for 17th IHIW principal investigators and lab members, project leaders, and component affiliates and chairs. When laboratory-initiated subject IDs are submitted to the WS Database, those IDs are anonymized and linked to unique 17th IHIW IDs, which are used to identify those subjects in genotyping and analysis efforts, to avoid the distribution of protected health information. The WS Database also stores project-specific data, using custom document formats, and analytic results.

#### 2.2.1 Participant Initiated Management of Typing Reports

The submission and management of typing reports is illustrated in Figure 1. Genotyping data and meta-data can be manually entered into the IHIW Database, and laboratory-generated and Illumina IHIW XML typing reports can be submitted directly to the IHIW Database. HML and GenDx XML typing reports must be converted to IHIW XML reports by uploading them to the sFTP server, and using the IHIW Database tools to generate IHIW XML reports from them. These converted IHIW XML typing reports are stored in the WS Database, where they are available for download by participants. Regardless of their source, all IHIW XML typing reports are submitted to and stored in the IHIW Database. Detailed instructions on the 17th IHIW data-submission process are available online^D^.

**Figure 1.**
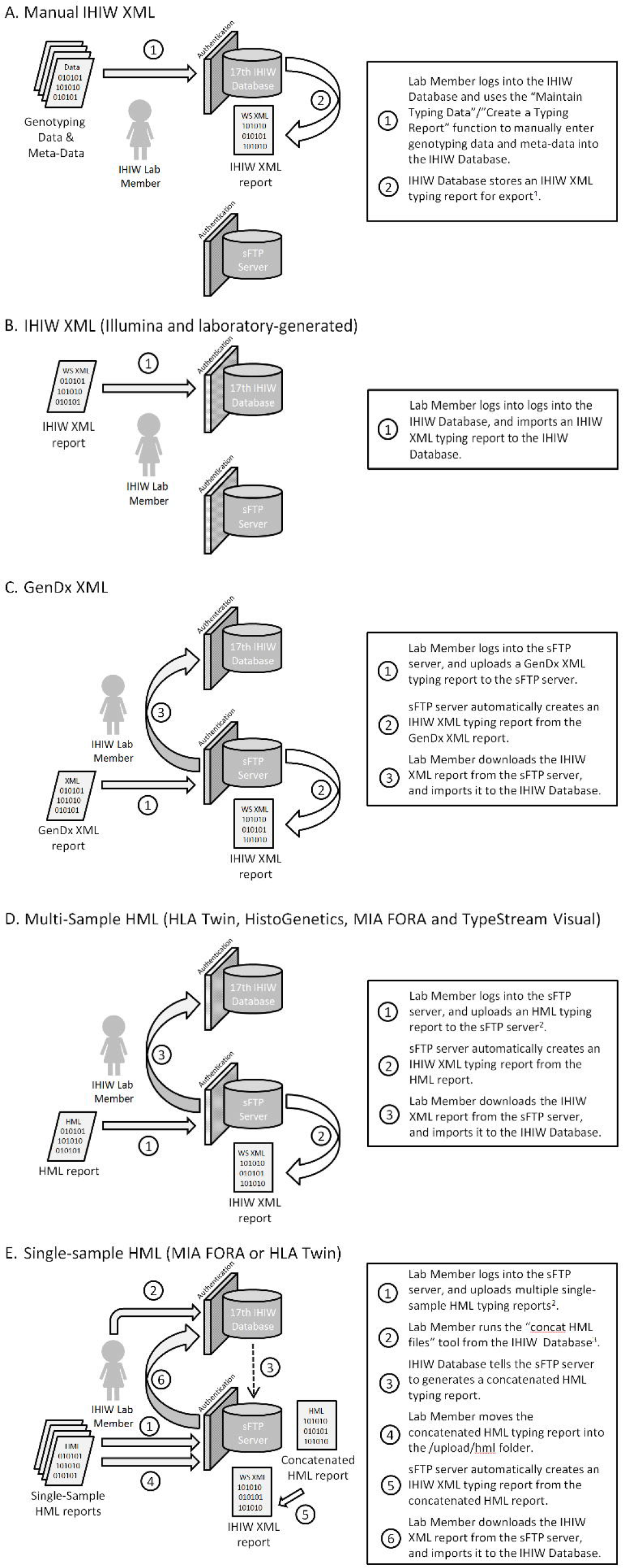
Typing Report Submission Procedures.

### 2.3 17th IHIW Standards and Tools

To facilitate uniform NGS data collection for the 17th IHIW, we have adopted specific data-standards and conventions for the validation of typing reports, and the analysis of workshop data. The tools described in sections 2.3.1 to 2.3.4 are available on GitHub^E,F^.

#### 2.3.1 Typing Report Validation

Given the number of typing report formats accepted by the WS Database, we have developed a number of tools and services for validating the format and content of each. Several of these tools are built into the WS Database, and run when typing reports are uploaded or created in the system. The semantic validations and WS Database functions applied to each typing report format are listed in Table 2. Because HML and GenDx XML typing reports are converted into IHIW XML reports, the validation and functions listed for IHIW WS format are applied to all typing reports. In addition, software developers generating HML typing reports have been encouraged to use the public MIRING validator for HML service (miring-validator^E^)) as part of their development efforts. This validator determines if a potential HML typing report follows basic HML and MIRING rules of syntax, and if it contains MIRING data-elements. Because this validator operates as a web-service, it can be built into an HML typing report generation pipeline.

**Table 2.** Validation and Database Functions Applied to each typing report format.

|  |  |  |
| --- | --- | --- |
| HML | Validation | 1. GL-String content validation with strict-mode GL service of the provided Reference Database version<br>2. GL-String “sanity check” syntax validation |
|  | Functions | 1. GL-String LiftOver to Reference Database version 3.25.0<br>2. GL-String concatenation<br>3. Retrieve HLA typing from GL-String using the provided reference allele |
| GenDx | Functions | 1. Generate GL-String from GenDx “genotype list” elements<br>2. Generate phasing groups |
| IHIW XML/Manual Entry | Validation | 1. Identifier (e.g. 17th IHIW labcode, sample ID) validation<br>2. GL-String validation with strict-mode 3.25.0 GL service<br>3. The uniqueness of the quartet annotation (sample ID, HLA typing, phasing group, start position) of consensus sequences<br>4. IUPAC nucleotide code[50] validation for consensus sequence |
|  | Functions | 1. hlaPoly application to identify novel polymorphisms |

HML: Histoimmunogenetics Markup Language
IHIW: International HLA and Immunogenetics Workshop
IUPAC: International Union of Pure and Applied Chemistry
XML: eXtensible Markup Language

#### 2.3.2 IPD-IMGT/HLA Database Versions

As noted in section 1.2.1, the Reference Database is updated quarterly; the number of alleles increases with each release, and the extent of sequence known for a given allele, as well as the number of fields in a given allele name, can increase between database releases. We address this by “freezing” all WS Database functions at Reference Database version 3.25.0. While HML or GenDx XML can be used to submit HLA allele names described in other Reference Database versions, the WS Database will translate those names to their 3.25.0 counterparts upon submission (as described in section 2.3.4), and all HLA-related data will be analyzed using Reference Database version 3.25.0, which is the source of all reference allele sequences. Restricting the WS Database to a single Reference Database version in this way streamlines database functions, facilitates uniform data management and analysis, and allows the final WS Database product to be updated to later Reference Database versions in future workshops. The Reference Database resources described below are available from the Reference Database FTP site^G^.

To facilitate the use of Reference Database 3.25.0 for the 17th IHIW, we have defined a set of full-length (“genomic”) version 3.25.0 reference alleles (Table 3) for use in generating and aligning consensus sequences, and identifying novel polymorphism. Though some alleles in this set may have names with fewer than four fields, indicating that no synonymous or non-coding polymorphism has been identified for those alleles as of Reference Database version 3.25.0, genomic sequence is available for all of them. When possible, a reference allele has been identified for each allele family at a locus, but for some loci a single full-length reference allele is identified.

**Table 3.** Full-length HLA Reference Alleles in IPD-IMGT/HLA Database Version 3.25.0.

| Locus | Accession Number | Allele Name | Description <sup>a</sup> |
| --- | --- | --- | --- |
| HLA-A | HLA00001 | HLA-A*01:01:01:01 | HLA-A Reference<br>A*01 Reference<br>A*36 Reference |
|  | HLA00005 | HLA-A*02:01:01:01 | A*02 Reference |
|  | HLA00037 | HLA-A*03:01:01:01 | A*03 Reference |
|  | HLA00043 | HLA-A*11:01:01:01 | A*11 Reference |
|  | HLA00048 | HLA-A*23:01:01 | A*23 Reference |
|  | HLA00050 | HLA-A*24:02:01:01 | A*24 Reference |
|  | HLA00071 | HLA-A*25:01:01 | A*25 Reference<br>A*26 Reference |
|  | HLA00085 | HLA-A*29:01:01:01 | A*29 Reference |
|  | HLA00089 | HLA-A*30:01:01 | A*30 Reference |
|  | HLA00097 | HLA-A*31:01:02:01 | A*31 Reference |
|  | HLA00101 | HLA-A*32:01:01 | A*32 Reference |
|  | HLA00104 | HLA-A*33:01:01 | A*33 Reference |
|  | HLA00108 | HLA-A*34:01:01 | A*34 Reference |
|  | HLA00112 | HLA-A*66:01:01 | A*43 Reference<br>A*66 Reference |
|  | HLA05918 | HLA-A*68:01:01:02 | A*68 Reference<br>A*69 Reference |
|  | HLA05527 | HLA-A*74:02:01:02 | A*74 Reference |
|  | HLA00130 | HLA-A*80:01:01:01 | A*80 Reference |
| HLA-B | HLA00132 | HLA-B*07:02:01 | HLA-B Reference<br>B*07 Reference<br>B*82 Reference<br>B*83 Reference |
|  | HLA00146 | HLA-B*08:01:01:01 | B*08 Reference |
|  | HLA00152 | HLA-B*13:01:01 | B*13 Reference |
|  | HLA00157 | HLA-B*14:01:01 | B*14 Reference |
|  | HLA00162 | HLA-B*15:01:01:01 | B*15 Reference |
|  | HLA00213 | HLA-B*18:01:01:01 | B*18 Reference |
|  | HLA00221 | HLA-B*27:02:01 | B*27 Reference |
|  | HLA00237 | HLA-B*35:01:01:01 | B*35 Reference |
|  | HLA00265 | HLA-B*37:01:01 | B*37 Reference |
|  | HLA00267 | HLA-B*38:01:01 | B*38 Reference<br>B*39 Reference |
|  | HLA00292 | HLA-B*40:01:02 | B*40 Reference |
|  | HLA13397 | HLA-B*40:305 | B*41 Reference |
|  | HLA00315 | HLA-B*42:01:01 | B*42 Reference |
|  | HLA00318 | HLA-B*44:02:01:01 | B*44 Reference |
|  | HLA00329 | HLA-B*45:01:01 | B*45 Reference |
|  | HLA00331 | HLA-B*46:01:01 | B*46 Reference |
|  | HLA14088 | HLA-B*47:01:01:03 | B*47 Reference |
|  | HLA00335 | HLA-B*48:01:01 | B*48 Reference<br>B*81 Reference |
|  | HLA00340 | HLA-B*49:01:01 | B*49 Reference<br>B*50 Reference |
|  | HLA00344 | HLA-B*51:01:01:01 | B*51 Reference |
|  |  |  | B*52 Reference<br>B*78 Reference |
|  | HLA00364 | HLA-B*53:01:01 | B*53 Reference<br>B*58 Reference |
|  | HLA00367 | HLA-B*54:01:01 | B*54 Reference<br>B*55 Reference<br>B*56 Reference<br>B*59 Reference |
|  | HLA00381 | HLA-B*57:01:01 | B*57 Reference |
|  | HLA00390 | HLA-B*67:01:01 | B*67 Reference |
|  | HLA00392 | HLA-B*73:01 | B*73 Reference |
| HLA-C | HLA00401 | HLA-C*01:02:01 | HLA-C Reference<br>C*01 Reference |
|  | HLA00405 | HLA-C*02:02:02:01 | C*02 Reference |
|  | HLA01543 | HLA-C*03:02:02:01 | C*03 Reference |
|  | HLA00420 | HLA-C*04:01:01:01 | C*04 Reference |
|  | HLA00427 | HLA-C*05:01:01:01 | C*05 Reference |
|  | HLA00430 | HLA-C*06:02:01:01 | C*06 Reference |
|  | HLA00433 | HLA-C*07:01:01:01 | C*07 Reference |
|  | HLA00445 | HLA-C*08:01:01 | C*08 Reference |
|  | HLA00454 | HLA-C*12:02:02 | C*12 Reference |
|  | HLA00462 | HLA-C*14:02:01:01 | C*14 Reference |
|  | HLA00467 | HLA-C*15:02:01:01 | C*15 Reference |
|  | HLA00475 | HLA-C*16:01:01:01 | C*16 Reference |
|  | HLA00481 | HLA-C*17:01:01:01 | C*17 Reference |
|  | HLA00483 | HLA-C*18:01 | C*18 Reference |
| HLA-DPA1 | HLA06604 | HLA-DPA1*01:03:01:02 | HLA-DPA1 Reference<br>DPA1*01 Reference<br>DPA1*03 Reference<br>DPA1*04 Reference |
|  | HLA00505 | HLA-DPA1*02:01:02 | DPA1*02 Reference |
| HLA-DPB1 | HLA00517 | HLA-DPB1*02:01:02 | HLA-DPB1 Reference |
| HLA-DQA1 | HLA00601 | HLA-DQA1*01:01:01:01 | HLA-DQA1 Reference |
| HLA-DQB1 | HLA00622 | HLA-DQB1*02:01:01 | HLA-DQB1 Reference |
| HLA-DRB1 | HLA00664 | HLA-DRB1*01:01:01 | HLA-DRB1 Reference<br>DRB1*01 Reference<br>DRB1*10 Reference |
|  | HLA00671 | HLA-DRB1*03:01:01:01 | DRB1*03 Reference |
|  | HLA00685 | HLA-DRB1*04:01:01:01 | DRB1*04 Reference |
|  | HLA00719 | HLA-DRB1*07:01:01:01 | DRB1*07 Reference |
|  | HLA00727 | HLA-DRB1*08:03:02 | DRB1*08 Reference |
|  | HLA09928 | HLA-DRB1*09:21 | DRB1*09 Reference |
|  | HLA00751 | HLA-DRB1*11:01:01:01 | DRB1*11 Reference |
|  | HLA14829 | HLA-DRB1*12:01:01:02 | DRB1*12 Reference |
|  | HLA00797 | HLA-DRB1*13:01:01:01 | DRB1*13 Reference |
|  | HLA00837 | HLA-DRB1*14:05:01 | DRB1*14 Reference |
| HLA-DRB1 | HLA03453 | HLA-DRB1*15:01:01:02 | DRB1*15 Reference<br>DRB1*16 Reference |
| HLA-DRB3 | HLA00887 | HLA-DRB3*01:01:02:01 | HLA-DRB3 Reference |
| HLA-DRB4 | HLA00905 | HLA-DRB4*01:01:01:01 | HLA-DRB4 Reference |
| HLA-DRB5 | HLA00915 | HLA-DRB5*01:01:01 | HLA-DRB5 Reference |

#### 2.3.3 Genotype Format and Validation

The large variety of formats used in the H&I community to record HLA genotypes and describe typing ambiguity makes it difficult to collect genotyping data in a uniform manner. We address this by collecting all HLA genotypes in GL string format[40], using the strict-mode GL Service[41] to validate the allele content of the GL string, and applying python scripts (pyglstring^E^) to validate the structure of the GL string. Data submitters are notified when GL strings fail validation (see section 2.3.5.2.1), and are requested to modify them accordingly.

These structural validation scripts include an exception for the DRB3, DRB4 and DRB5 loci (the secondary DRB loci), permitting combinations of alleles at these loci to be connected by the GL string “+” operator (e.g., “HLA-DRB3*01:26N+HLA-DRB5*01:01:01”), whereas for other loci, the “+” operator connects only alleles of a single locus. When GL string format was introduced[40], the “+” operator denoted the number of copies of a given gene present in an individual. Ideally, given the structural haplotype variation known for the DRB loci[46], when the absence of a DRB3, DRB4 or DRB5 locus can be determined, the absence of that locus should be noted in a GL string. The WMDA guidelines indicate that the absence of any allele at a secondary DRB locus be reported using “NNNN” (e.g., “HLA-DRB1*NNNN”)[38], but this is not a widely used approach. Without a standard nomenclature for describing the confirmed absence of a secondary DRB gene, we treat these loci as alleles of a single locus. The development of a nomenclature for describing the confirmed absence of a locus (e.g. “HLA-DRB3*NNNN”, “HLA-DRB3*00:00” or “HLA-DRB3*ABSENT”) should be considered by the Nomenclature Committee.

#### 2.3.4 LiftOver Tool

As typing reports are accepted into the WS Database, HLA genotypes identified under Reference Database versions other than 3.25.0 are translated to their 3.25.0 counterparts via a LiftOver tool (IHIW17LiftOver.pm^F^). Non-3.25.0 alleles are translated on the basis of their Reference Database accession numbers, as related in the Allelelist_history.txt file^G^. In cases of alleles named after version 3.25.0 (e.g., HLA-A*01:01:01:05, identified in Reference Database version 3.27.0), the submitted allele name is translated to either the lowest-numbered 3.25.0 allele name with the greatest number of matching lower-order fields to the submitted allele (e.g., HLA-A*01:01:01:01 is chosen to replace HLA-A*01:01:01:05), or the reference allele for that locus (Table 3) when there are no matching lower-order fields, and the submitted allele is noted in the “Novelpolymorphism” field for that genotype (e.g. as “IPD-IMGT/HLA-3270-HLA-A*01:01:01:05”). In cases where allele name changes that occurred in Reference Database versions prior to 3.25.0 resulted in accession number changes (e.g., HLA-DRB1*08:01:03, with accession number HLA02257, was changed to HLA-DRB1*08:01:01, with accession number HLA00723, as part of Reference Database version 3.24.0, as detailed in Table 4), the version 3.25.0 allele name is used.

**Table 4.** HLA Allele Remapping for WS Database LiftOver and Consensus Linking Functions.

| Reference Database Release in which the change occurred | Rationale | Original Allele Name | Original Accession Number | Current Allele Name | Current Accession Number | Current Allele Present in Version 3.25.0? | Accession Number Change? | WS Database Action When Original Allele Name Has Been Submitted |
| --- | --- | --- | --- | --- | --- | --- | --- | --- |
| 3.17.0 | Sequence identical | B*49:15 | HLA05834 | B*49:01:01 | HLA00340 | YES | YES | Use 3.25.0 version of Current Allele Name |
| 3.20.0 | Sequence renamed | A*26:03:02 | HLA04741 | A*26:111 | HLA04741 | YES | NO |  |
| 3.21.0 | Sequence error | DRB1*11:11:02 | HLA02157 | DRB1*11:11:01 | HLA00765 | YES | YES |  |
| 3.22.0 | Sequence error | A*03:194 | HLA11939 | A*03:213 | HLA12966 | YES | YES |  |
| 3.24.0 | Sequence error | A*23:69 | HLA12676 | A*23:01:01:01 | HLA00048 | YES<sup>1</sup> | YES |  |
|  | Sequence identical | DRB1*08:01:03 | HLA02257 | DRB1*08:01:01 | HLA00723 | YES | YES |  |
| 3.25.0 | Sequence renamed | DRB1*04:94:02N | HLA14178 | DRB1*04:212N | HLA14178 | YES | NO |  |
| 3.26.0 | Sequence renamed | A*30:02:12 | HLA09547 | A*30:100 | HLA14873 | YES | YES | Use 3.25.0 version of Original Allele Name |
|  | Sequence error | C*17:01:01:01 | HLA00481 | C*17:01:01:02 | HLA04311 | YES | YES |  |
| 3.27.0 | Sequence renamed | DPB1*35:01:02 | HLA04110 | DPB1*621:01 | HLA04110 | NO<sup>2</sup> | NO |  |
|  | Sequence identical | DQB1*06:220 | NA<sup>3</sup> | DQB1*06:217 | HLA16016 | NO<sup>2</sup> | NA<sup>3</sup> | Change to DQB1*06:01:01<sup>4</sup> |
| 3.28.0 | Sequence identical | DPA1*02:02:01 | HLA00508 | DPA1*02:07:01:01 | HLA15619 | NO<sup>5</sup> | YES | Use 3.25.0 version of Original Allele Name |
| 3.29.0 | Sequence identical | DQB1*03:01:01:13 | HLA07476 | DQB1*03:01:01:07 | HLA17167 | NO<sup>6</sup> | YES | Change to DQB1*03:01:01:01<sup>7</sup> |

When ambiguous HLA genotypes are submitted, the LiftOver tool evaluates ambiguous alleles (delimited with the GL string slash [/] operator) and ambiguous genotypes (delimited with the GL string pipe [|] operator), and identifies alleles and genotypes that can be translated to their 3.25.0 counterparts (illustrated in Figure 2). These alleles are translated, and the GL string is consolidated to eliminate duplications. If an ambiguous HLA genotype consists entirely of alleles that were named after Reference Database version 3.25.0, the LiftOver tool translates those alleles to the corresponding lowest-numbered 3.25.0 alleles with the greatest number of matching lower-order fields, as described above, and consolidates the GL string. In all cases, the submitted non-3.25.0 GL strings are stored in the “Original_GL” field for that genotype. These allelic and GL string LiftOver functions are accomplished using a modified version of the Allelelist_history.txt file that includes data from the hla_nom.txt^G^ files and Table 3 (IHIW17_AllelelistGgroups_history.txt^F^). This LiftOver process occurs when HML and GenDx XML typing reports are converted to IHIW XML reports. All IHIW XML typing reports correspond to version 3.25.0.

**Figure 2.**
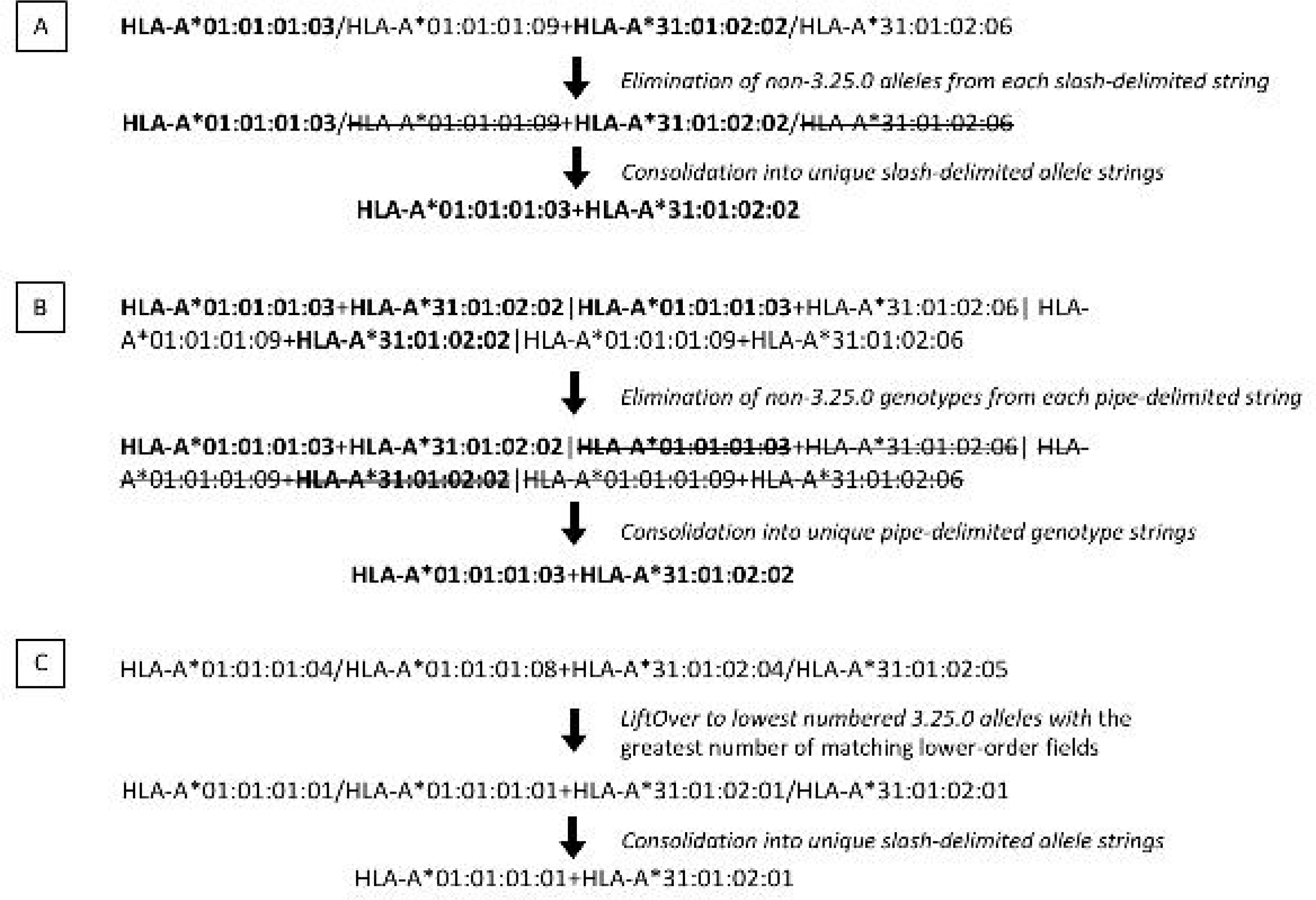
Example of LiftOver of Ambiguous HLA-A Genotypes.

#### 2.3.5 17th IHIW Database Tools and Functions

Reference Database version 3.25.0 includes 12.9 million bases of sequence for 14,957 HLA alleles at 19 HLA loci. Of this, more than 40,000 exons comprise 9 million bases of sequence, making this a rich, but very complex, data resource. We are developing user-facing front end tools to assist 17th IHIW participants in working with these data, and data-facing back end tools to facilitate the integration of the large quantities of new sequence that will be generated via NGS.

##### 2.3.5.1 Front End Tools

###### 2.3.5.1.1 hlaPoly

The absence of a standard method for describing novel nucleotide polymorphism in consensus sequences poses challenges for our uniform data collection approach. Typing reports generated for the same specimen using different genotyping software may include identical consensus sequences and genotypes, but when a consensus sequence includes novel polymorphism, the Reference Database version, reference allele sequence, and sequence coordinate system used to describe that polymorphism can vary between software applications, and typing reports generated by different software may identify different novel polymorphism for identical consensus sequences. For example, the nucleotide sequence of the HLA-A*01:01:01:05 allele differs from the HLA-A*01:01:01:01 allele in Reference Database 3.25.0 at three intron 2 nucleotide positions, and differs from the HLA-A*01:01:01:03 allele in Reference Database 3.25.0 at those same three positions as well as at an intron 1 position; the reference allele used to align the HLA-A*01:01:01:05 consensus sequence informs the description of novel polymorphism.

To standardize novel polymorphism description for the 17^th^ IHIW, we developed the hlaPoly R package^F^, which identifies novel polymorphism for a given consensus sequence, when provided with the closest matching allele name (which is usually what is included in the genotype) and the Reference Database version (currently, version 3.25.0). The hlaPoly tool is deployed online as a Shiny application^H^. As illustrated in Supplementary Figure S2, hlaPoly uses the DECIPHER R package[47] to generate a multiple sequence alignment for the full-length HLA reference allele sequence (Table 3), the sequence of the pertinent allele in the genotype (closest allele) and the consensus sequence, and then retrieves the mismatches and indels between the consensus sequence and the called allele as novel polymorphism. If no sequence is known for the called allele in an aligned region, the mismatches and indels between the consensus sequence and the full-length HLA reference allele are retrieved. For each novel polymorphism, the feature number and start/end position relative to that feature are also calculated. The WS Database stores these novel polymorphism data in both a tabular form (see the bottom of Figure S1) and a string format (described in Supplement A).

###### 2.3.5.1.2 Quick Calculation of Feature Position

To assist in manual entry of genotyping data and meta-data into the WS Database, we have developed a tool for the calculation of gene-feature information. Given an allele name and the nucleotide position relative to start of known nucleotide sequence for that allele, the tool returns the feature name (e.g. Exon 2), feature ID and the relative nucleotide position in that feature. This tool is available in the WS Database under “Lab Member”/”Tools”/”IMGT/HLA Feature List”.

###### 2.3.5.1.4 Concatenate HML files

Each HML file uploaded to the sFTP server is treated as single typing report, and as suggested in Figure 1, some HML typing reports are generated for individual samples. Rather than requiring that hundreds or thousands of individual-sample HML files be converted to IHIW XML files, each of which would need to be manually loaded into the WS Database, we have provided a tool (concathml.pl^F^)that concatenates multiple HML files into a single HML file, which can be converted into a single IHIW XML file for loading. This “Concat HML files” tool available in the WS Database under “Lab Member”/”Tools”.

###### 2.3.5.1.4 “Convert HML to IHIW XML” and “Convert GenDX XML to IHIW XML”

As noted in Figure 1, the sFTP server will automatically generate an IHIW XML typing report when an HML is loaded into the /hml directory. The server will also generate an IHIW XML report when an GenDX XML report is loaded into the /gendx directory. The “Convert HML to IHIW XML” and “Convert GenDx XML to IHIW XML” tools can be used to force these automatic functions to run immediately, or to manually convert HML or GenDx XML typing reports loaded into other directories. Both tools are available in the WS Database under “Lab/Member”/”Tools”.

##### 2.3.5.2 Back End Tools

###### 2.3.5.2.1 Watcher Daemons

To monitor activity on the sFTP server, we have developed daemons that detect new HML and GenDx XML files as they are uploaded to the sFTP server, automatically convert them to IHIW XML files, and validate them during the conversion. Any validation errors are logged and made available under “Lab Member”/”Tools”/”Job Log” in the WS Database. A second set of daemons perform daily checks for new typing reports in the WS Database. These daemons run hlaPoly for newly added or edited consensus sequences, and store the novel polymorphism results in the WS Database.

###### 2.3.5.2.2 Consensus Linking

Genotypes and consensus sequences are recorded separately in HML typing reports. Each consensus sequence is associated with the reference allele used align it, which is usually a full-length allele, but is not directly linked to specific alleles in the associated genotype. For cases when these reference alleles are not included in the genotype, we have developed a consensus linking tool that identifies the allele in the genotype that most closely matches the reference allele using the same approach applied for the LiftOver process (described in section 2.3.4). For example, if HLA-A*11:01:01:01 and *31:01:02:01 are the respective reference alleles for consensus sequences A and B, which are associated with the HLA-A*11:01:28+HLA-A*31:01:07, the consensus linking tool would associate HLA-A*11:01:28 with consensus sequence A and *31:01:07 with consensus sequence B.

### 2.4 Support for 17th IHIW Projects

In addition to its collection, validation and storage functions, the WS Database supports 17th IHIW projects by integrating tools for HLA data analysis and exchange. An updated version of PyPop[48]^I^, supporting colon-delimited allele names, with increased multi-locus analysis capacity, will be accessible through the WS Database system. Similarly, integration of Gene Feature Enumeration[49] (GFE) functions (e.g., the feature-service^E,J^, GFE service^E,K^ and Allele-Calling Tool^E,L^) into the WS Database system will allow full-gene HLA sequences to be exchanged and analyzed in the absence of an HLA allele name.

## 3. Conclusions

We have addressed several of the long-standing challenges to uniform NGS HLA data-collection and - storage by developing new tools and formats, and adopting existing standards and services. NGS vendors have worked with us to develop equivalent NGS HLA typing reports that ensure data-portability across 17th IHIW projects. We ensure data-quality by validating all typing reports before they are loaded to the WS Database. All HLA genotyping data are recorded using the same Reference Database version, and novel HLA polymorphism is described using the same reference alleles. This approach will facilitate the basic and clinical research aims of 17^th^ IHIW Projects, and the larger H&I community. The 17^th^ IHIW will be held in September of 2017. Our ultimate goal is for the WS Database to serve as a central H&I community resource that will persist after 17^th^ IHIW and ensure research and data continuity with future IHIW efforts.

## Acknowledgements

This work was supported National Institutes of Health (NIH) National Institute of Allergy and Infectious Disease (NIAID) grant R01AI128775 (BM, MM, SM), NIH National Institute of General Medical Sciences (NIGMS) grant R01GM109030 (BM, MM, SM), Office of Naval Research (ONR) grant N00014-08-1-1207 (BM and MM), and an overseas project funded by the Taiwanese Ministry of Science and Technology (MST) (CC). The content is solely the responsibility of the authors and does not necessarily reflect the official views of the MST, NIAID, NIGMS, NIH, ONR, Taiwanese government or United States government. We thank the Stanford Blood Center for the support and promotion of the 17th IHIWS endeavor, Ken Yamaguchi for helpful discussions and manuscript review, Tamara Vayntrub for her tremendous administrative support of 17th IHIW efforts, the histocompatibility and immunogenetics community and the International HLA and Immunogenetics Workshop Council for their continued dedication to and support of the International Workshops, and President Barack H. Obama for his support and appreciation of American science and basic research.

